# Longitudinal White Matter Microstructure and Transition to Psychosis in Individuals at Clinical High Risk

**DOI:** 10.64898/2026.09.27.754817

**Authors:** Lara Bolte, George Gifford, Mauricio Serpa, Michiel Cottaar, Paola Dazzan, Paolo Fusar-Poli, Stefania Tognin, Alexis E. Cullen, Matthew J. Kempton, Margot I. E. Slot, Hendrika van Hell, Arija Maat, Lieuwe de Haan, Benedicto Crespo Facorro, Birte Glenthøj, Stephen Lawrie, Thérèse van Amelsvoort, Celso Arango, Irina Falkenberg, Barnaby Nelson, Silvana Galderisi, Rodrigo A. Bressan, Jun Soo Kwon, Kang Ik K. Cho, Mark Weiser, Romina Mizrahi, Gabriele Sachs, Matthias Kirschner, Maxime Taquet, PSYSCAN Consortium, Rene Kahn, Dominic OIiver, Philip McGuire

## Abstract

Altered white matter microstructure, commonly indexed by fractional anisotropy (FA), has been reported in individuals at clinical high-risk of psychosis (CHR-P) across multiple white matter tracts. However, the relationship between these alterations and later transition to psychosis remains unclear. To address this gap, we investigated whether baseline and longitudinally updated FA measurements were associated with risk of transition to psychosis in a large cohort of individuals at CHR-P.

We analysed diffusion tensor imaging data from *N* = 205 participants at CHR-P, 23 of whom developed psychosis within 3 years of follow-up. Data were collected at baseline, 6 months, and 12 months. Associations between global and tract-specific FA and transition risk were examined using Cox proportional hazards models of baseline data and longitudinally updated data.

Point estimates were directionally consistent, with hazard ratios <1 for FA across all white matter tracts in both baseline and longitudinal models. Associations reached nominal significance in the hippocampal cingulum, anterior thalamic radiation, and forceps minor, but none of these remained significant after correcting for multiple comparisons (*p*_corr_ > .05).

The directionally consistent estimates are compatible with small, distributed effects across multiple white matter pathways being associated with transition to psychosis, although this requires replication. Longitudinally updated FA measurements did not yield stronger associations than were available from baseline imaging data alone. Studies at an even larger scale are needed to establish whether these associations are robust and clinically meaningful.

## 1. Introduction

Meta-analyses suggest that approximately 20-25% of individuals meeting clinical high-risk for psychosis (CHR-P) criteria will transition to a first-episode psychosis (FEP) within 2 to 3 years ^1,2^. Identifying biological markers associated with transition may improve our understanding of the mechanisms that underlie psychosis onset and, ultimately, contribute to more accurate risk stratification.

Psychosis is thought to involve alterations in white matter (WM) microstructure, most commonly assessed using fractional anisotropy (FA), a metric derived from diffusion tensor imaging (DTI) ^3^. Changes in FA have also been identified in individuals at CHR-P and individuals with FEP, with cross-sectional studies predominantly reporting widespread reductions in FA in both groups compared with healthy controls ^4–8^. While heterogeneous and few in number, longitudinal studies indicate that WM microstructure may continue to change during the CHR-P stage ^9^. Importantly, such differences in FA have been associated with clinical symptoms at baseline and longitudinally ^10,11^. Together, these findings suggest that WM alterations are already present before the onset of psychosis and may be clinically meaningful.

Evidence linking FA to subsequent transition is considerably more limited. Some studies have reported lower FA at baseline in individuals who later transition to psychosis compared with those who do not ^10,12,13^ and healthy controls ^10,12,14–16^, implicating widespread regions including frontotemporal association fibres, projection fibres, and callosal tracts. Additionally, reductions in global FA at baseline have been associated with transition after 12 months ^17^. However, others have reported no baseline differences in free-water adjusted FA between those who did or did not transition to psychosis ^18^, or even lower FA in non-transitioners ^16^, highlighting the inconsistency of the current evidence. Importantly, these findings are based on baseline imaging and therefore cannot determine whether transition to psychosis is related to within-subject changes in FA after baseline.

Longitudinal DTI evidence remains particularly sparse. Early studies reported progressive reductions in frontal WM microstructure in CHR-P and associations between changes in FA and subsequent clinical outcomes, but the sample sizes in these studies were small ^15,19^. More recent work on free-water adjusted FA in a larger multisite cohort found little evidence for an association between longitudinal changes in FA related to transition, instead suggesting subtle deviations in age-related WM development in CHR-P compared with healthy controls ^18^.

Overall, the relationship between longitudinal WM changes in individuals at CHR-P and transition to psychosis remains uncertain. In particular, it is unclear whether serial FA measurements during the CHR-P stage may provide additional information about transition risk beyond data from imaging at baseline. To address this question, the present study analysed data from CHR-P participants in the PSYSCAN study using a time-to-event approach incorporating time-varying FA. This allowed both baseline and updated measurements across multiple WM tracts to be examined in relation to subsequent transition risk. We collected DTI data from a large cohort of individuals at CHR-P across nine different sites, scanning participants again after 6 and 12 months, and including clinical follow-up for 3 years to determine which individuals developed psychosis.

We first tested the hypothesis that lower baseline global and tract-specific FA would be associated with an increased risk of subsequent transition to psychosis. We further hypothesised that incorporating serial FA measurements would provide additional information about transition risk beyond baseline FA alone.

## 2. Methods and Materials

### 2.1. Study Design and Participants

This study used data from the PSYSCAN study, an international, longitudinal, multisite cohort study conducted across nine sites: Australia (Melbourne), Italy (Naples), the Netherlands (Amsterdam, Maastricht), Spain (Madrid), United Kingdom (London), Canada (Toronto), South Korea (Seoul), Brazil (São Paulo). Recruitment took place between July 2016 and December 2019. Comprehensive details of the study protocol, recruitment procedures, and eligibility criteria have been published elsewhere ^20–22^.

Participants were individuals at CHR-P, defined using the Comprehensive Assessment of At-Risk Mental States (CAARMS)^23^ or the Schizophrenia Proneness Instrument, Adult version (SPI-A)^24^. Exclusion criteria included (i) previous neurosurgery or neurological disorder, (ii) IQ < 70, (iii) inability to provide informed consent, (iv) head injuries resulting in loss of consciousness, (v) ages outside the eligible recruitment range (16-40 years; 14-40 at Madrid), (vi) magnetic resonance imaging (MRI) contraindications (including pregnancy), (vii) more than 30 days cumulative exposure to antipsychotic medication at a dosage for treating first-episode psychosis in the 3 months prior to baseline, and (viii) any past episode of frank psychosis lasting more than 7 days.

Ethical approval was obtained at each participating site, and all participants, or their legal representatives for participants < 18 years old, provided written informed consent in accordance with the Declaration of Helsinki.

### 2.2. Diffusion Tensor Imaging

Diffusion MRI acquisition, preprocessing, and site harmonisation procedures were identical to those described previously ^25^. Briefly, imaging was performed at all sites using harmonised single-shell, echo-planar, diffusion-weighted protocols on 3 Tesla MRI scanners. Preprocessing included brain extraction (SynthStrip) ^26,27^, correction for susceptibility-induced distortions (Synb0-DISCO ^28,29^ and TOPUP ^30,31^), and motion and eddy-current correction (EDDY) ^32^, followed by quality control performed using QUAD/SQUAD ^33^.

FA maps were generated using DTIFIT^31^ and processed with tract-based spatial statistics (TBSS)^34^. Global FA was calculated, together with FA values from 20 major WM tracts defined using the Johns Hopkins University probabilistic WM atlas^35^. To reduce statistical comparisons, the left and right homologous tracts were averaged, and the temporal and non-temporal components of the superior longitudinal fasciculus were combined, resulting in 11 regions of interest: (1) anterior thalamic radiation, (2) corticospinal tract, (3) cingulum – cingulate gyrus, (4) cingulum – hippocampus, (5) forceps major, (6) forceps minor, (7) inferior fronto-occipital fasciculus, (8) inferior longitudinal fasciculus, (9) superior longitudinal fasciculus, (10) uncinate fasciculus, (11) global FA (Figure 1).

**Figure 1.**
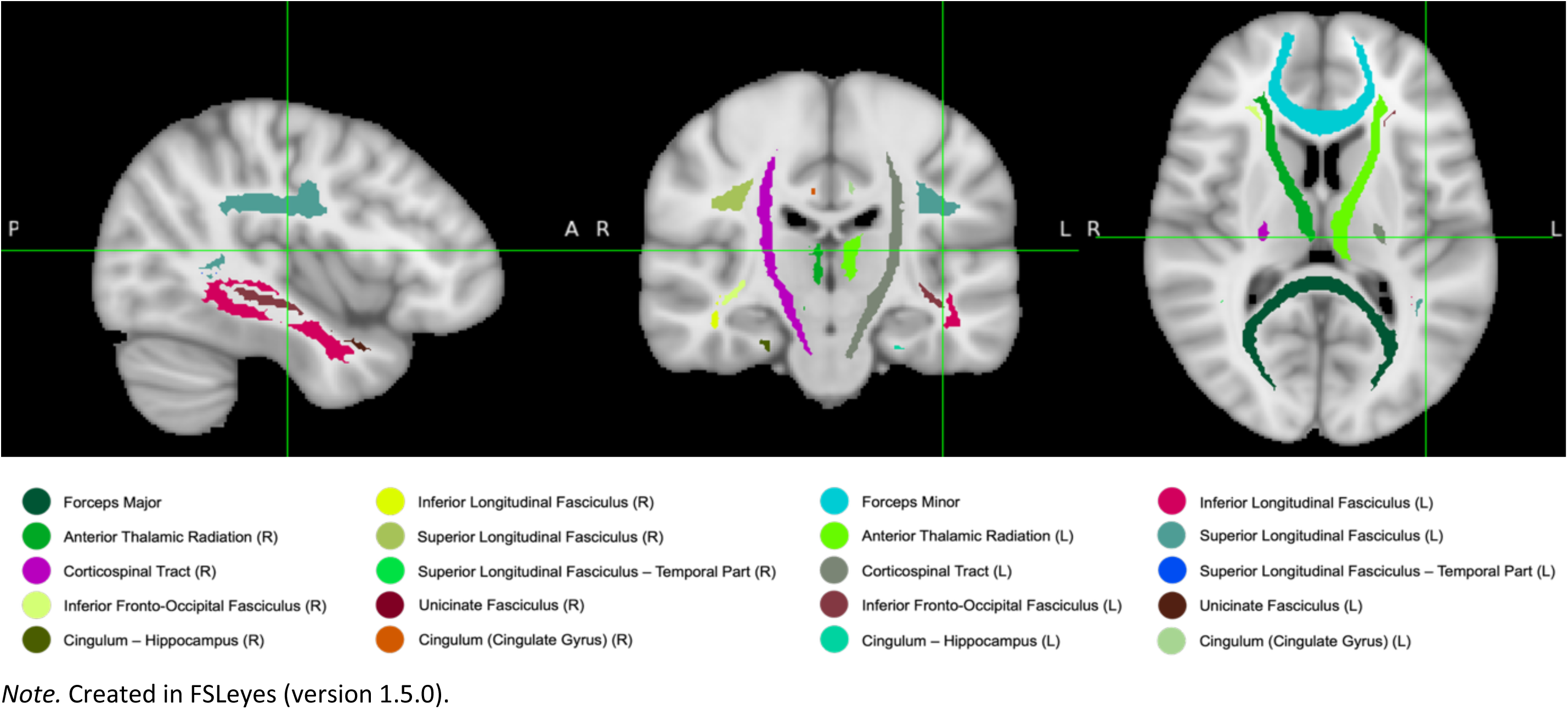
White Matter Tracts (JHU Atlas)

To minimise scanner-related variability across sites, diffusion measures were harmonised using the travelling-heads mean-offset correction and longitudinal ComBat (LongCombat)^36^, adjusting for demographic, clinical, and imaging covariates, as described previously ^25^.

### 2.3. Transition Status

Transition to psychosis was defined during study clinical assessments using CAARMS criteria. For participants in whom transition occurred between study visits, or after study completion or withdrawal, transition was determined using clinical data from electronic health records (EHR), with transition defined as any non-organic psychotic disorder according to the Diagnostic and Statistical Manual of Mental Disorders (DSM) or the International Classification of Diseases (ICD), or by accepted referrals to early intervention for psychosis services. When data from both a study clinical assessment and an EHR-defined transition were available, the time of transition was defined by the earliest recorded date of the transition. For participants who did not transition, follow-up time was censored at the date of the last available clinical assessment or EHR confirmation of non-transition.

For the primary analyses, survival was administratively censored at 3 years (1095.75 days) to increase stability of survival estimates, resulting in two transitions occurring after this cut-off being treated as censored observations and recoded as non-transitions. This censoring cut-off was selected because previous studies have shown that the majority of transitions to psychosis occur within the first 3 years, after which risk plateaus ^1^. Additionally, this approach reduced the temporal gap between the final neuroimaging scan (12 months) and potential transition date, thereby improving the temporal correspondence between FA measurements and the survival outcome.

### 2.4. Statistical Analyses

#### 2.4.1. Data Preparation

Analyses were conducted in R (version 4.5.2) and focused on FA measures, including global FA and 10 tract-specific FA values. Prior to model fitting, several data preparation steps were undertaken.

To ensure stable model estimation, ethnicity was collapsed from seven categories into three groups based on the largest number of participants: White, East Asian, and Other. This was necessary because several of the original categories contained very few or no transition events, which would have resulted in unstable parameter estimates in the Cox models.

As Cox proportional hazards models require complete covariate data, missing baseline covariate values were handled using multiple imputation by chained equations, implemented with the *mice* package (version 3.19.0). Missingness was observed for gender (*n* = 1; 0.49%), ethnicity (*n* = 2; 0.98%), chlorpromazine (CPZ) equivalent dose at baseline (*n* = 16; 7.80%), and WAIS IQ (*n* = 20; 9.76%), whereas age data were complete. Twenty imputed datasets were generated. Predictive mean matching was used for continuous variables (CPZ equivalent dose and WAIS IQ), logistic regression for the binary variable (gender), and multinomial logistic regression for the categorical variable (ethnicity). Time-to-transition and transition status were excluded from the imputation model, and the imputed baseline covariates were subsequently carried forward across all longitudinal survival intervals.

All FA measures were z-standardised before modelling. Consequently, hazard ratios (HRs) represent the change in hazard associated with a 1 SD increase in FA relative to the distribution of the respective FA measure in the analysis sample.

### 2.4.2. Survival Analysis

To test whether baseline FA was associated with transition to psychosis, separate Cox proportional hazards models were fitted using the *survival* package (version 3.8.3) for global FA and each tract-specific FA measure (11 measures total), examining whether baseline FA was associated with transition to psychosis. All models were adjusted for age, gender, ethnicity, CPZ equivalent dose, and IQ, and used the Efron approximation to handle tied event times.

To test the second hypothesis, namely whether incorporating longitudinally updated FA measurements was associated with transition risk, we fitted extended Cox proportional hazards models with time-varying covariates for global FA and each tract-specific FA measure. Unlike the baseline model, which assumes that baseline FA remains representative throughout follow-up, this approach allows FA values to be updated when subsequent imaging measurements become available, thereby incorporating within-participant changes in FA over time when estimating transition risk. Data were structured using the counting-process (start-stop) formulation, whereby each participant contributed one or more observation intervals defined by consecutive assessment times. Within each interval, the most recently acquired FA measurement was used as the time-varying variable and carried forward until the next FA measurement, occurrence of the event, or end of follow-up. As with the baseline models, the longitudinal Cox models were adjusted for age, gender, ethnicity, CPZ equivalent dose, and IQ, with cluster-robust standard errors used to account for repeated observations within participants. Similarly, the Efron approximation was used to handle tied event times.

Both the baseline and longitudinal analyses were conducted separately within each of the 20 imputed datasets, and parameter estimates were combined using Rubin’s rules. To account for multiple testing across the 11 FA-specific analyses, *p*-values were adjusted using the Benjamini-Hochberg false discovery rate (FDR) procedure. Statistical significance was defined as a two-sided FDR-adjusted *p* < .05. Results are reported as HRs with 95% confidence intervals (CIs). The proportional hazards assumption was assessed using Schoenfeld residuals on one completed imputed dataset, as no standard pooling procedure exists for Schoenfeld residual tests following multiple imputation.

#### 2.4.3. Robustness Analyses

Several analyses were conducted to evaluate the robustness of the primary findings. First, to assess whether the findings were robust to the bilateral averaging of homologous WM tracts used in the primary analyses, the Cox models were repeated using left-and right-hemispheric FA measures separately. This resulted in 21 FA measures: anterior thalamic radiation (L/R), corticospinal tract (L/R), cingulum – cingulate gyrus (L/R), cingulum – hippocampus (L/R), forceps major, forceps minor, inferior fronto-occipital fasciculus (L/R), inferior longitudinal fasciculus (L/R), superior longitudinal fasciculus (L/R), superior longitudinal fasciculus – temporal part (L/R), uncinate fasciculus (L/R), global FA.

Second, previous studies have shown that individuals meeting Brief Limited Intermittent Psychotic Symptoms (BLIPS) criteria have a higher risk of transition to psychosis than non-BLIPS individuals ^1^. We therefore examined whether the association between FA and transition risk was modified by BLIPS status using interaction terms in the Cox models. A fully subgroup-stratified analysis was not performed because of the limited number of transition events and substantial overlap between subgroups.

Third, given evidence suggesting higher transition rates in male individuals at CHR-P than in females ^37^, we examined whether the association between FA and transition risk was modified by gender using interaction terms in the Cox models.

Finally, to evaluate the impact of the predefined follow-up limit, the primary analyses were repeated without censoring follow-up at 3 years.

## 3. Results

### 3.1. Participant Characteristics

DTI data were available for *N* = 219 participants. Of these, *n* = 14 participants were excluded from the analysis because of missing clinical follow-up data, which precluded construction of survival intervals. The final analysis cohort thus comprised 205 participants, of whom *n* = 23 (11%) transitioned to psychosis within the 3-year follow-up period (DTI: 23 at BL, 12 at 6M, 9 at 12M; 69.6% men; mean age 21.5±4.1 years; Table 1) and *n* = 182 did not (DTI: 176 at BL, 113 at 6M, 108 at 12M; 52.2% men; mean age 22.9±4.8 years). Overall, 19/205 (9.3%) met BLIPS criteria, including 5/23 (21.7%) of those who transitioned.

**Table 1.** Sociodemographic Characteristics.

|  | No Transition | Transition |
| --- | --- | --- |
| Number of Participants |  |  |
| Total *N* | 182 | 23 |
| Baseline | 176 | 23 |
| 6-months | 112 | 13 |
| 12-months | 107 | 10 |
| Sociodemographic Variables |  |  |
| Age in years |  |  |
| Mean (*SD*) | 22.9 (4.8) | 21.5 (4.1) |
| Gender |  |  |
| Male *n* (%) | 95 (52.2%) | 16 (69.6%) |
| Ethnicity |  |  |
| White | 109 (59.9%) | 11 (47.8%) |
| East Asian | 25 (13.7%) | 8 (34.8%) |
| Other Ethnic Minority | 46 (25.3%) | 4 (17.4%) |
| WAIS IQ |  |  |
| Mean (*SD*) | 105.8 (18.2) | 102.2 (14.4) |
| Antipsychotic dosage mg/day |  |  |
| Mean (*SD*) | 17.2 (72.2) | 17.1 (32.6) |
| Clinical Variables |  |  |
| CAARMS |  |  |
| Positive Symptom Severity: Mean (*SD*) | 11.0 (3.8) | 12.7 (3.8) |
| Negative Symptom Severity: Mean (*SD*) | 6.0 (3.6) | 5.6 (3.1) |
| SOFAS |  |  |
| Mean (*SD*) | 53.3 (11.1) | 54.3 (12.0) |

*Note.* Sociodemographic and clinical variables were assessed at baseline.

None of the 23 who transitioned to psychosis met the criteria for frank psychosis at baseline, consistent with the study inclusion criteria. The cumulative number of transitions progressively increased over follow-up, with 4 participants having transitioned by 6 months, 10 by 12 months, 16 by 18 months, 19 by 24 months, 22 by 30 months, and 23 by 36 months (Figure 2).

**Figure 2.**
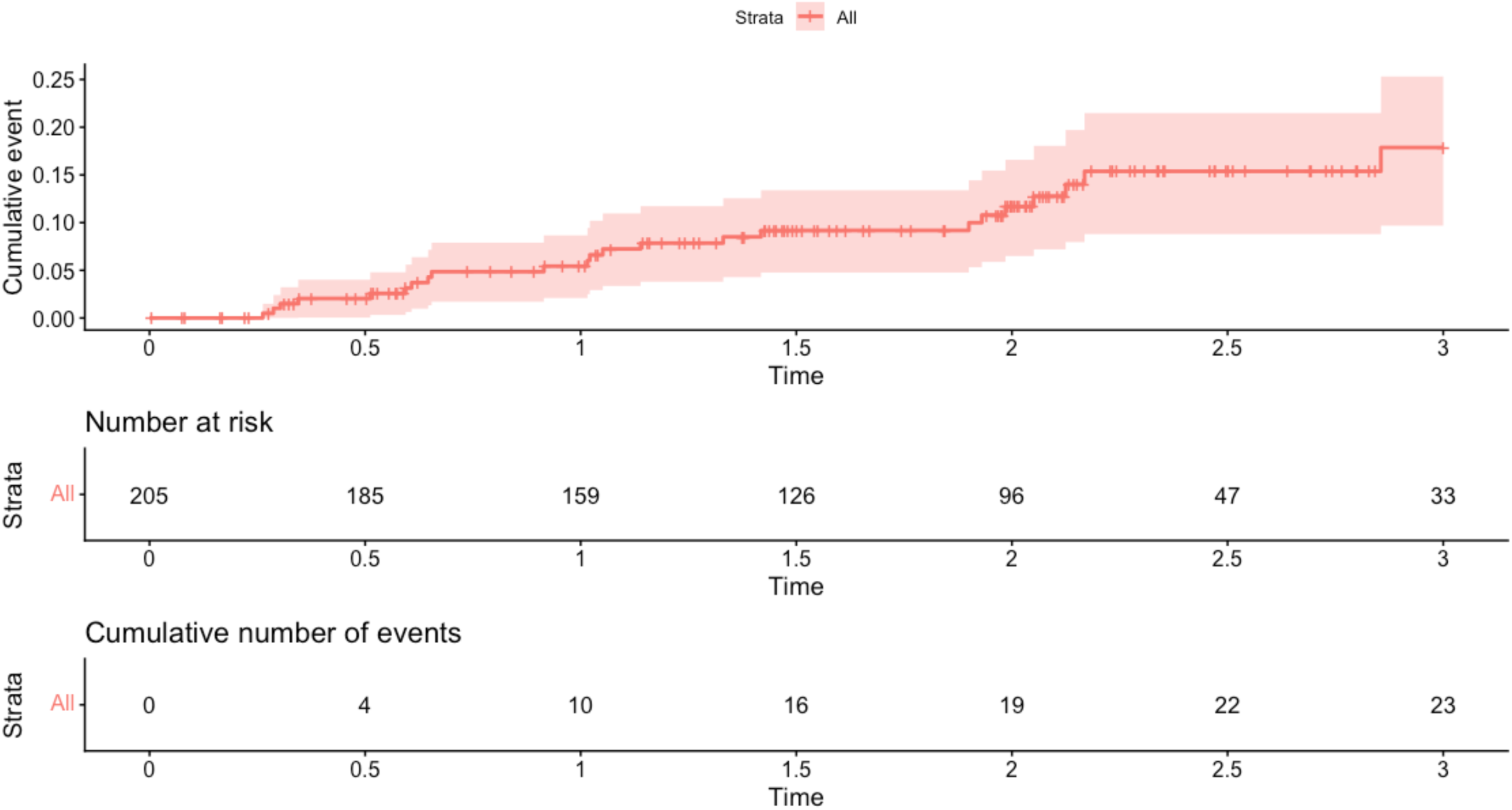
Kaplan-Meier Plot Illustrating Cumulative Incidence of Transition to Psychosis.

The mean follow-up time was 644 days (SD = 322, median = 722, IQR = 384-860). Among participants who transitioned to psychosis within the 3 years, the mean number of days from baseline to transition was 431 days (SD = 272; median = 373, IQR = 220-700). The 3-year cumulative incidence of psychosis was 17.87% (95% CI: 9.67%-25.31%).

### 3.2. Survival Analyses

Across both the baseline and longitudinal Cox proportional hazards models, higher FA was consistently associated with a lower hazard of transition to psychosis. HRs for a 1 SD increase in FA were below 1.0 for global FA and for all 10 tract-specific measures, ranging from 0.50 to 0.75 (95% CI 0.28–1.20) in the baseline models and from 0.48 to 0.62 (95% CI 0.23–1.46) in the longitudinal models (Table 2), albeit with wide 95% CIs.

**Table 2.** Hazard Ratios Consistently <1 Across All White Matter Tracts.

| Tract | Baseline Model |  |  | Longitudinal Model |  |  |
| --- | --- | --- | --- | --- | --- | --- |
| | HR (95% CI) | $p$ | $p_{\text{corr}}$ | HR (95% CI) | $p$ | $p_{\text{corr}}$ |
| Global FA | 0.64 (0.39 – 1.04) | .069 | .165 | 0.51 (0.24 – 1.09) | .073 | .150 |
| Anterior Thalamic Radiation | 0.70 (0.44 – 1.12) | .125 | .165 | 0.48 (0.23 – 0.97) | *.043 | .150 |
| Cingulum Cingulate Gyrus | 0.69 (0.43 – 1.11) | .118 | .165 | 0.62 (0.32 – 1.23) | .137 | .150 |
| Cingulum Hippocampus | 0.50 (0.28 – 0.90) | *.023 | .165 | 0.59 (0.24 – 1.46) | .197 | .197 |
| Corticospinal Tract | 0.70 (0.44 – 1.13) | .135 | .165 | 0.61 (0.31 – 1.22) | .130 | .150 |
| Forceps Major | 0.68 (0.43 – 1.09) | .100 | .165 | 0.52 (0.27 – 1.00) | .051 | .150 |
| Forceps Minor | 0.75 (0.47 – 1.20) | .215 | .215 | 0.53 (0.28 – 1.00) | *.049 | .150 |
| Inferior Fronto Occipital Fasciculus | 0.69 (0.43 – 1.11) | .116 | .165 | 0.53 (0.26 – 1.05) | .064 | .150 |
| Inferior Longitudinal Fasciculus | 0.66 (0.40 – 1.07) | .087 | .165 | 0.54 (0.24 – 1.21) | .110 | .150 |
| Superior Longitudinal Fasciculus | 0.70 (0.43 – 1.13) | .133 | .165 | 0.60 (0.29 – 1.24) | .134 | .150 |
| Uncinate Fasciculus | 0.73 (0.46 – 1.16) | .166 | .183 | 0.55 (0.27 – 1.13) | .089 | .150 |

*Note.* HRs represent the change in hazard of transition to psychosis per 1-SD higher FA; $p_{\text{corr}}$ = $p$ -value FDR-corrected for multiple comparisons.
\* $p < .05$

In the baseline analyses, lower FA in the hippocampal part of the cingulum (HR = 0.50, 95% CI 0.28–0.90, *p* = .022) was associated with an increased hazard of later transition to psychosis.

In the longitudinal analyses, lower FA in the anterior thalamic radiation (HR = 0.48, 95% CI 0.24–0.97, *p* = .043) and forceps minor (HR = 0.53, 95% CI 0.28–1.00, *p* = .049) were similarly associated with an increased hazard of transition to psychosis (Figures 3 and 4).

**Figure 3.**
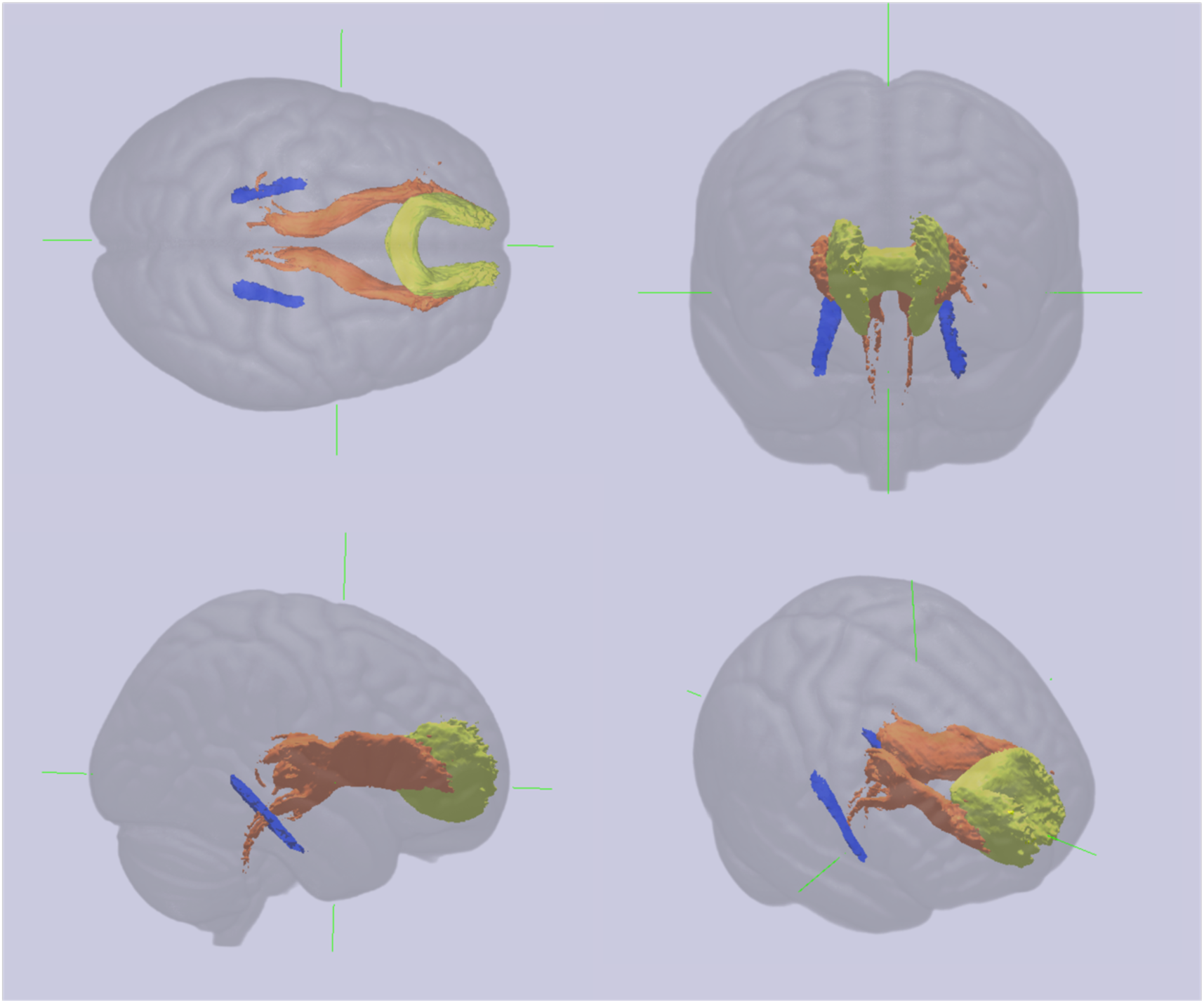
White Matter Tracts with Nominal Associations with Transition to Psychosis Risk.

**Figure 4.**
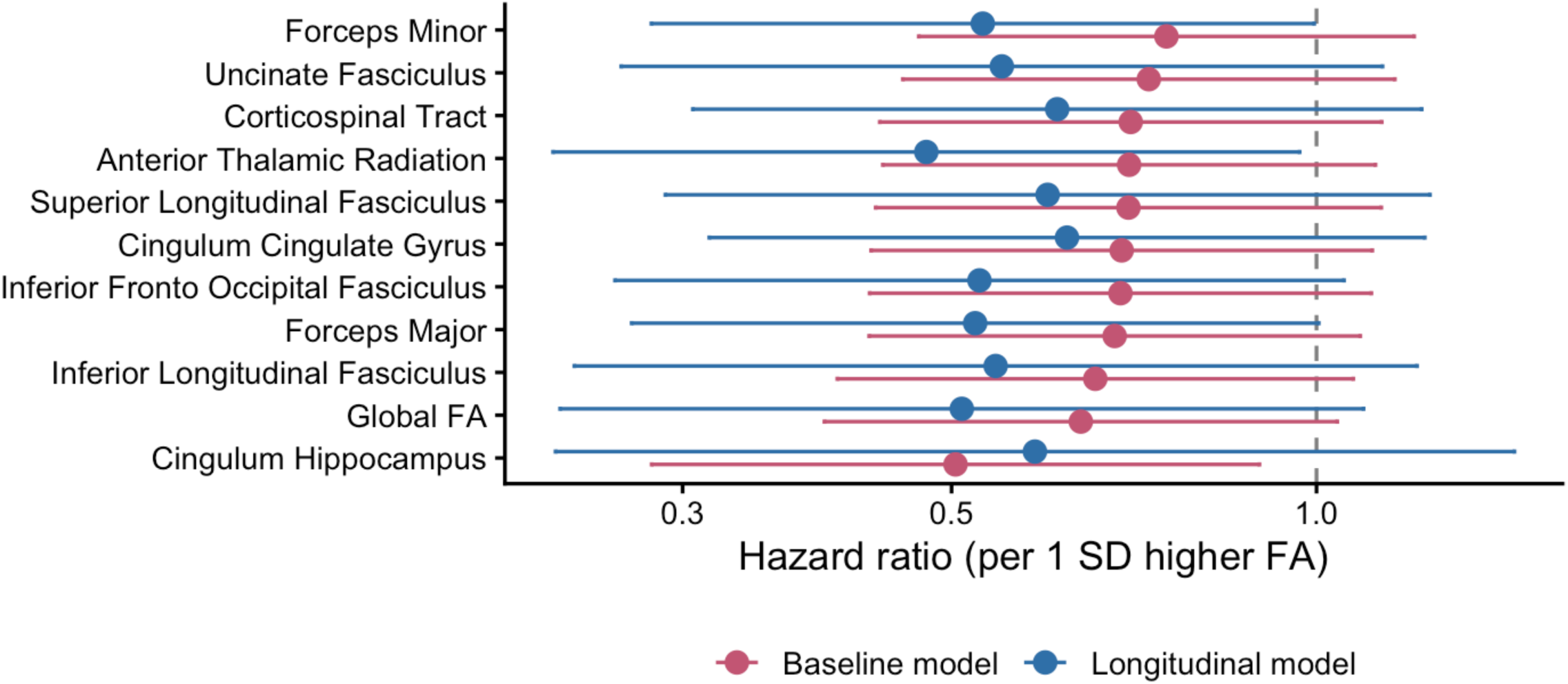
Baseline and Longitudinal Cox Proportional Hazards Models.

No other tracts were significantly associated with transition to psychosis across baseline and longitudinal models (*p* > .05). After correction for multiple comparisons, none of the associations between FA and transition to psychosis described above remained statistically significant (all FDR-adjusted *p*_corr_ > .175 for baseline models and *p*_corr_ > .150 for longitudinal models). There was no evidence of violation of the proportional hazards assumption for any predictor or for the global test in either the baseline or longitudinal Cox models (all *p* > .05).

### 3.3. Robustness Analyses

All sensitivity analyses produced findings consistent with the primary Cox proportional hazards models, with no association surviving correction for multiple comparisons.

Analysing WM tracts for each hemisphere separately yielded results that were consistent with the primary analyses (Table S1; Figure S1). HRs remained below 1 for all tract measures and global FA, ranging from 0.51 to 0.76 in the baseline models and from 0.48 to 0.70 in the longitudinal models. The smallest nominal *p*-value in the baseline analysis was observed for the left cingulum hippocampus (HR = 0.51, 95% CI 0.29–0.89, *p* = .022) and right cingulum hippocampus (HR = 0.56, 95% CI 0.33–0.95, *p* = .035), whereas the longitudinal analysis showed the strongest nominal associations for the right anterior thalamic radiation (HR = 0.48, 95% CI 0.23–0.98, *p* = .045), left anterior thalamic radiation (HR = 0.48, 95% CI 0.24–0.98, *p* = .046), and forceps minor (HR = 0.53, 95% CI 0.28–1.00, *p* = .049). However, no association remained significant after FDR correction (all FDR-adjusted *p*_corr_ ≥ .158).

Including an interaction between FA and BLIPS status did not provide evidence that the association between FA and transition was related to BLIPS status (Table S2). Although several interaction terms reached nominal significance in the baseline models (smallest *p* = .011 for forceps major), none survived FDR correction, and no nominally significant interactions were observed in the longitudinal analyses. Interaction term HRs were predominantly greater than 1 in the baseline models, whereas the longitudinal models showed a more mixed pattern of effect estimates.

There was no evidence that gender modified the association between FA and transition risk (Table S3). No FA × gender interaction reached nominal significance in either the baseline (all *p* ≥ .56) or longitudinal analyses (all *p* ≥ .14), and all remained non-significant following FDR correction.

Repeating the analyses without censoring follow-up at 3 years produced results that were highly consistent with the primary analyses (Table S4). HRs remained below 1 across all FA measures. Although two additional tract-specific associations reached nominal significance in the longitudinal analyses, namely the forceps major and the inferior fronto-occipital fasciculus, none survived FDR correction.

## 4. Discussion

This study investigated the association between WM microstructure and risk of transition to psychosis in individuals at CHR-P. In line with the hypothesised direction of effect, HR estimates consistently indicated a higher hazard of transition with lower FA across all WM tracts, both at baseline and when incorporating longitudinally updated FA measurements. Several tract-specific associations reached nominal statistical significance. However, none survived correction for multiple comparisons.

Despite the lack of statistically significant findings after multiple comparison correction, the consistent direction of effect across all examined tracts is notable. The tracts showing the strongest evidence of an association between transition risk and FA were the hippocampal cingulum (a limbic pathway with connections to the medial temporal cortex ^38,39^), the anterior thalamic radiation (a projection pathway linking the thalamus with the prefrontal cortex ^39^) and the forceps minor (a callosal tract connecting the prefrontal regions across the two hemispheres ^40^). This overall pattern is broadly in line with previous evidence suggesting that WM abnormalities associated with transition to psychosis are widespread, involving frontotemporal association pathways ^10,14,19^, callosal tracts ^13–16^, projection fibres ^12^, and limbic regions ^10^. The anatomical distribution of the nominal findings is also consistent with evidence from neuroimaging and preclinical systematic reviews implicating these tracts and their connected brain regions in psychosis more broadly ^4,41,42^, providing some face validity for the observed findings despite their lack of statistical significance after correction. Together, these observations are compatible with a small, distributed relationship between WM microstructure and psychosis onset rather than a strong effect confined to a single anatomical pathway. However, because FA values are correlated across tracts, the consistency of the HRs should not be interpreted as independent evidence for a common global effect.

Importantly, incorporating longitudinally updated FA measurements produced slightly stronger associations for several tracts, particularly in the anterior thalamic radiation and forceps minor. However, contrary to the second hypothesis, it did not substantially alter the overall pattern of findings seen with the baseline data alone. The longitudinal models were also associated with wider confidence intervals than the baseline models, potentially due to the greater complexity of the longitudinal models and attrition of subjects with imaging data over time. This is broadly consistent with previous longitudinal studies, which reported progressive reductions in WM microstructure associated with transition while also observing similar associations in baseline analyses ^15,19^. A more recent study found little evidence that either baseline or longitudinal free-water adjusted FA distinguished individuals who transitioned from those who did not ^18^. Unlike these previous studies, which used repeated-measures ANOVA ^15,19^ or linear mixed-effects models ^18^, the present study’s time-to-event framework allowed repeated imaging assessments to contribute dynamically to estimation of transition risk. Despite this different analytical approach, longitudinally updated FA measurements provided limited additional information about transition risk beyond baseline FA within the temporal resolution of the present cohort.

The absence of statistically significant associations after multiple-comparison correction should also be considered in the context of the relatively small number of transition events. Possible underlying associations are likely to be modest and distributed across multiple WM tracts, making adequate statistical power particularly challenging to achieve. Because only a minority of individuals at CHR-P transition to psychosis, with further reductions in available data due to attrition over follow-up, relatively large baseline samples are needed to yield a sufficient number of transition events. In the present study, 23 of the 205 participants (11%) transitioned within 3 years, substantially fewer than the approximately 20-25% transition rate typically reported in CHR-P cohorts ^1,2^. Consequently, the present study may have been underpowered to detect relatively modest associations between WM microstructure and transition risk. This interpretation is consistent with the data from the sensitivity analyses, in which the 3-year censoring limit modestly strengthened the observed associations, while increasing the number of comparisons through hemispheric-specific analyses modestly weakened the associations, albeit with limited changes in statistical significance.

There was no evidence that the association between FA and transition differed according to BLIPS status or gender. However, these interaction analyses should be interpreted cautiously because relatively few transition events occurred within the relevant subgroups, resulting in imprecise estimates with wide confidence intervals. Consequently, modest subgroup-specific effects cannot be excluded.

Overall, these findings suggest that FA alone is unlikely to provide a strong marker of transition to psychosis. However, the consistency in the direction of effects across WM pathways raises the possibility that small, distributed associations may contribute alongside other clinical and neurobiological factors.

### 4.1. Limitations and Future Directions

Several limitations should be acknowledged. First, despite the relatively large multisite CHR-P cohort, the number of participants who later transitioned to psychosis was modest, limiting statistical power to detect small associations. This may have been complicated by the inherent clinical heterogeneity of CHR-P populations, which may increase variability and obscure neurobiological associations with subsequent transition. Future large, harmonised multisite CHR-P cohorts with greater numbers of transition events will be important for determining whether the small, distributed effects observed here are robust and clinically meaningful. Second, ethnicity was collapsed into broad categories, limiting the extent to which ethnic variation could be examined within PSYSCAN’s internationally diverse cohort. Future studies could also complement categorical transition outcomes with continuous measures of psychosis risk or clinical progression. Capturing variation across the full CHR-P cohort may provide greater sensitivity to WM characteristics associated with progression towards psychosis and help determine whether the distributed associations observed here extend beyond the subgroup who transition within a defined follow-up period. Third, although FA is a widely used diffusion metric in studies of psychosis, it is a non-specific measure of WM microstructure and cannot determine which biological processes have contributed to the altered signal. Thus, it is unclear if the results are related to differences in myelination, axonal density, fibre coherence, or crossing fibres. Where feasible, advanced diffusion acquisition protocols, such as multi-shell imaging, should be considered in future studies to enable more biologically informative modelling approaches, e.g. NODDI ^43^ or BENCH ^44^. This may provide greater insight into the microstructural alterations associated with transition to psychosis.

### 4.2. Conclusions

In this prospective, multisite longitudinal study, lower FA was consistently associated with a higher estimated hazard of transition across widespread WM tracts at the level of point estimates, although no association survived correction for multiple comparisons. Incorporating longitudinally updated FA provided little additional information beyond baseline measurements. Together, these findings suggest that any association between FA and transition to psychosis is likely to be modest and distributed and requires replication in larger cohorts with greater numbers of transition events.

## Acknowledgments

Special thanks to Gaurav Bhalerao and Jacob Turnbull for their expertise in site harmonisation and Jesper Anderson for his guidance during the preprocessing of the diffusion data.

## 5. Sources of Funding

The PSYSCAN Project is supported by grant agreement n° 603196 under the European Union’s Seventh Framework Programme. This research is partly supported by the National Institute for Health Research (NIHR) Mental Health Biomedical Research Centre (BRC) at South London and Maudsley NHS Foundation Trust.

LB, GG, MT, DO, and PM are supported by the NIHR Oxford Health BRC. The views expressed are those of the author(s) and not necessarily those of the NHS, the NIHR or the Department of Health.

CA was supported by the Spanish Ministry of Science and Innovation, Instituto de Salud Carlos III (ISCIII), co-financed by the European Union, ERDF Funds from the European Commission, “A way of making Europe”, financed by the European Union – NextGenerationEU (PMP21/00051), PI19/01024. PI22/01824 CIBERSAM, Madrid Regional Government (B2017/BMD-3740 AGES-CM-2), European Union Structural Funds, European Union Seventh Framework Program, European Union H2020 Program under the Innovative Medicines Initiative 2 Joint Undertaking: Project PRISM-2 (Grant agreement No.101034377), Project AIMS-2-TRIALS (Grant agreement No 777394), Horizon Europe, the National Institute of Mental Health of the National Institutes of Health under Award Number 1U01MH124639-01 (Project ProNET) and Award Number 5P50MH115846-03 (project FEP-CAUSAL), Fundación Familia Alonso, and Fundación Alicia Koplowitz.

## 6. Collaborators

Kate Merritt ^1^, Natalia Petros ^1^, Mathilde Antoniades ^1^, Andrea De Micheli ^1^, Sandra Vieira ^1^, Tom Spencer ^1^, Gemma Modinos ^2^, Frederike Schirmbeck ^3^, Diana Tordesillas-Gutierrez ^4,5^, Esther Setien-Suero ^4,5^, Rosa Avesa-Arriola ^4,5^, Paula Suarez-Pinilla ^4,5^, Victor Ortiz Garcia-de la Foz ^4,5^, Mikkel Erlang Sørensen ^6^, Bjørn H. Ebdrup ^6,7^, Karen Tangmose ^6,7^, Helle Schæbel ^6^, Egill Rostrup ^6,8^, Oliver Gruber ^9^, Anja Richter ^1,9^, Bernd Krämer ^9^, Bea Campforts ^10^, Machteld Marcelis ^10^, Claudia Vingerhoets ^10^, Covandonga M. Díaz-Caneja ^11^, Miriam Avora ^11^, Joost Janssen ^11^, Roberto Rodríguez-Jiménez ^12,13^, Marina Díaz-Marsá ^13^, Tilo Kircher ^14^, Florian Bitsch ^14^, Jens Sommer ^14^, Patrick McGorry ^15,16^, Paul Amminger ^15,16^, Meredith McHugh ^15,16^, Armida Mucci ^17^, Paola Bucci ^17^, Giuseppe Piegari ^17^, Daria Pietrafesa ^17^, Alessia Nicita ^17^, Sara Patriarca ^17^, André Zugman ^18^, Graccielle Rodrigues da Cunha ^18^, Tae Young Lee ^19^, Minah Kim ^19^, Sun Young Moon ^19^, Silvia Kyungjin Lho ^19^, Michael Kiang ^20,21,22^, Sarah Ahmed ^20,21^, Jenny Lepock ^20,21^, Margaret Maheandiran ^21^, Ivana Prce ^21^, Cory Gerritsen ^21,23^, Matthäus Willeit ^24^, Marzena Lenczowski ^24^, Ulrich Sauerzopf ^24^, Ana Weidenauer ^24^, Julia Furtner ^25^, Anke Maatz ^26^, Achim Burrer ^26^, Philipp Stämpfli ^26^, Naemi Huber ^26^, Stefan Kaiser ^27,28^, Wolfram Kawohl ^29^, Inge Winter-van Rossum ^30^

1. Department of Psychosis Studies, Institute of Psychiatry, Psychology & Neuroscience, King’s College London, London, UK

2. Department of Psychological Medicine, Institute of Psychiatry, Psychology & Neuroscience, King’s College London, London, UK

3. Amsterdam UMC, University of Amsterdam, Psychiatry, Department Early Psychosis, Amsterdam, The Netherlands

4. Department of Psychiatry, Marqués de Valdecilla University Hospital, IDIVAL. School of Medicine, University of Cantabria, Santander, Spain

5. CIBERSAM, Centro Investigación Biomédica en Red Salud Mental, Spain

6. Centre for Neuropsychiatric Schizophrenia Research (CNSR) & Centre for Clinical Intervention and Neuropsychiatric Schizophrenia Research (CINS), Mental Health Centre Glostrup, University of Copenhagen, Glostrup, Denmark

7. University of Copenhagen, Faculty of Health and Medical Sciences, Deptartment of Clinical Medicine, Copenhagen, Denmark

8. Functional Imaging Unit (FIUNIT), Rigshospitalet Glostrup, University of Copenhagen, Glostrup, Denmark

9. Section for Experimental Psychopathology and Neuroimaging, Department of General Psychiatry, Heidelberg University, Heidelberg, Germany

10. Department of Psychiatry and Neuropsychology, Maastricht University, Maastricht, The Netherlands

11. Servicio de Psiquiatría del Niño y del Adolescente, Hospital General Universitario Gregorio Marañon, Universidad Complutense Madrid, Spain; Centro de Investigación Biomédica en Red de Salud Mental, Madrid, Spain

12. Departmento de Psiquiatría, Instituto de Investigación Sanitaria Hospital 12 de Octubre (imas12), Madrid, Spain

13. Hospital Clínico de San Carlos, Universidad Complutense, Centro de Investigación Biomédica en Red de Salud Mental (CIBERSAM), Madrid, Spain

14. Department of Psychiatry, University of Marburg, Marburg, Germany

15. Orygen, The National Centre of Excellence in Youth Mental Health, 35 Poplar Road, Parkville, Victoria, Melbourne, Australia

16. Centre for Youth Mental Health, The University of Melbourne, Parkville, Victoria, Australia

17. Department of Psychiatry, University of Campania Luigi Vanvitelli, Largo Madonna delle Grazie, Naples, Italy

18. Department of Psychiatry, Interdisciplinary Lab for Clinical Neurosciences (LiNC), Universidade Federal de São Paulo (UNIFESP), São Paulo, Brazil

19. Department of Neuropsychiatry, Seoul National University College of Medicine, Jongno-gu, Seoul, South Korea

20. Institute of Medical Science, University of Toronto, Toronto, Ontario, Canada

21. Centre for Addiction and Mental Health, Toronto, Ontario, Canada

22. Department of Psychiatry, McGill University, Montreal, Canada

23. Department of Psychology, University of Toronto, Toronto, Ontario, Canada

24. Department of Psychiatry and Psychotherapy, Medical University of Vienna, Vienna, Austria

25. Department of Biomedical Imaging and Image-guided Therapy, Medical University of Vienna, Vienna, Austria

26. Department of Psychiatry, Psychotherapy and Psychosomatics, Psychiatric Hospital, University of Zurich, Switzerland

27. Clinical and Experimental Psychopathology Laboratory, Faculty of Medicine, University of Geneva, Switzerland

28. Adult Psychiatry Division, Department of mental health and psychiatry, University Hospitals of Geneva, Switzerland

29. Clienia Schloessli, Private Psychiatric Hospital and Academic Teaching Hospital of the University of Zurich, Oetwil am See, Zurich, Switzerland

30. Department of Psychiatry, UMC Utrecht Brain Center, University Medical Center Utrecht, Utrecht, The Netherlands

## Disclosures

CA has been a consultant to or has received honoraria or grants from Abbot, Acadia, Ambrosetti, Angelini, Biogen, BMS, Boehringer, Carnot, Gedeon Richter, Janssen Cilag, Lundbeck, Medscape, Menarini, Minerva, Otsuka, Pfizer, Roche, Sage, Servier, Shire, Schering Plough, Sumitomo Dainippon Pharma, Sunovion, Takeda and Teva.

SG has been a consultant and/or advisor to or has received honoraria from Angelini, Boehringer-Ingelheim, Gedeon Richter-Recordati, Janssen, Lundbeck, Otsuka, ROVI and Bristol Myers Squibb.

GS has been a consultant and/or advisor to, or has received honoraria from Boehringer Ingelheim, Janssen-Cilag, Lundbeck, Mylan, Sanova, Schwabe, Recordati.

MK has received consulting fees from Otsuka for activities unrelated to the present study. DO has received consulting fees from Google DeepMind for activities unrelated to the present study.

MS received support from Lundbeck to attend a scientific conference unrelated to the present study.

