## Supplementary material for "Longitudinal White Matter Microstructure and Transition to Psychosis in Individuals at Clinical High Risk": Table S1

*Hazard Ratios Consistently <1 Across All White Matter Tracts – No Hemisphere-Averaging*

| Tract | Baseline Model | | | Longitudinal Model | | |
|---|---|---|---|---|---|---|
| | HR (95% CI) | *p* | $p_{corr}$ | HR (95% CI) | *p* | $p_{corr}$ |
| Global FA | 0.64 (0.39 – 1.04) | .069 | .202 | 0.51 (0.24 – 1.09) | .073 | .158 |
| Anterior Thalamic Radiation - Left | 0.69 (0.43 – 1.11) | .115 | .202 | 0.48 (0.24 – 0.98) | *.046 | .158 |
| Anterior Thalamic Radiation - Right | 0.72 (0.45 – 1.14) | .144 | .202 | 0.48 (0.23 – 0.98) | *.045 | .158 |
| Cingulum Cingulate Gyrus - Left | 0.73 (0.45 – 1.18) | .181 | .211 | 0.59 (0.29 – 1.18) | .112 | .158 |
| Cingulum Cingulate Gyrus - Right | 0.66 (0.41 – 1.06) | .084 | .202 | 0.68 (0.34 – 1.34) | .207 | .217 |
| Cingulum Hippocampus - Left | 0.51 (0.29 – 0.89) | *.022 | .202 | 0.51 (0.22 – 1.21) | .105 | .158 |
| Cingulum Hippocampus - Right | 0.56 (0.33 – 0.95) | *.035 | .202 | 0.70 (0.29 – 1.68) | .353 | .353 |
| Corticospinal Tract - Left | 0.73 (0.46 – 1.17) | .173 | .211 | 0.62 (0.30 – 1.30) | .164 | .199 |
| Corticospinal Tract - Right | 0.68 (0.41 – 1.11) | .111 | .202 | 0.61 (0.32 – 1.16) | .106 | .158 |
| Forceps Major | 0.68 (0.43 – 1.09) | .100 | .202 | 0.52 (0.27 – 1.00) | .051 | .158 |
| Forceps Minor | 0.75 (0.47 – 1.20) | .215 | .226 | 0.53 (0.28 – 1.00) | *.049 | .158 |
| Inferior Fronto Occipital Fasciculus - Left | 0.72 (0.44 – 1.15) | .155 | .204 | 0.55 (0.28 – 1.07) | .069 | .158 |
| Inferior Fronto Occipital Fasciculus - Right | 0.67 (0.42 – 1.08) | .092 | .202 | 0.51 (0.25 – 1.06) | .064 | .158 |
| Inferior Longitudinal Fasciculus - Left | 0.69 (0.42 – 1.11) | .114 | .202 | 0.60 (0.29 – 1.26) | .142 | .186 |
| Inferior Longitudinal Fasciculus - Right | 0.64 (0.39 – 1.05) | .075 | .202 | 0.50 (0.21 – 1.18) | .094 | .158 |
| Superior Longitudinal Fasciculus - Left | 0.69 (0.43 – 1.12) | .125 | .202 | 0.58 (0.29 – 1.16) | .102 | .158 |
| Superior Longitudinal Fasciculus - Right | 0.69 (0.43 – 1.11) | .115 | .202 | 0.64 (0.31 – 1.33) | .180 | .199 |
| Superior Longitudinal Fasciculus Temporal Part - Left | 0.70 (0.43 – 1.13) | .135 | .202 | 0.57 (0.27 – 1.20) | .113 | .158 |
| Superior Longitudinal Fasciculus Temporal Part - Right | 0.75 (0.46 – 1.21) | .212 | .226 | 0.63 (0.30 – 1.32) | .172 | .199 |
| Uncinate Fasciculus - Left | 0.76 (0.47 – 1.21) | .226 | .226 | 0.57 (0.29 – 1.12) | .086 | .158 |
| Uncinate Fasciculus - Right | 0.71 (0.44 – 1.12) | .130 | .202 | 0.55 (0.26 – 1.16) | .097 | .158 |

**Table S2**

*Influence of Interaction Terms of BLIPS and White Matter on Hazard Ratios for Transition to Psychosis Risk*

| Tract | Baseline Model | | | Longitudinal Model | | |
|---|---|---|---|---|---|---|
| | HR (95% CI) | *p* | $p_{corr}$ | HR (95% CI) | *p* | $p_{corr}$ |
| Global FA | 3.24 (0.59 – 17.76) | .158 | .194 | 1.41 (0.07 – 30.17) | .764 | .788 |
| Anterior Thalamic Radiation | 2.20 (0.40 – 12.13) | .334 | .344 | 0.72 (0.08 – 6.34) | .685 | .759 |
| Cingulum Cingulate Gyrus | 3.66 (0.73 – 18.30) | .105 | .134 | 1.11 (0.08 – 15.04) | .919 | .919 |
| Cingulum Hippocampus | 2.17 (0.42 – 11.24) | .327 | .344 | 3.77 (0.47 – 30.56) | .149 | .497 |
| Corticospinal Tract | 0.93 (0.23 – 3.82) | .916 | .916 | 0.48 (0.05 – 4.24) | .396 | .577 |
| Forceps Major | 5.86 (1.62 – 21.23) | *.011 | .088 | 3.00 (0.42 – 21.38) | .190 | .512 |
| Forceps Minor | 2.67 (0.43 – 16.61) | .265 | .292 | 0.56 (0.10 – 3.24) | .400 | .577 |
| Inferior Fronto Occipital Fasciculus | 7.41 (1.20 – 45.58) | .033 | .088 | 1.62 (0.07 – 39.94) | .690 | .759 |
| Inferior Longitudinal Fasciculus | 5.61 (1.07 – 29.57) | *.043 | .088 | 2.24 (0.14 – 35.68) | .455 | .577 |
| Superior Longitudinal Fasciculus | 4.56 (0.88 – 23.65) | .068 | .104 | 1.93 (0.11 – 32.76) | .546 | .648 |
| Uncinate Fasciculus | 3.80 (0.49 – 29.30) | .181 | .206 | 0.76 (0.07 – 7.86) | .751 | .788 |

**Table S3**

*Influence of Interaction Terms of Gender and White Matter on Hazard Ratios for Transition to Psychosis Risk*

| Tract | Baseline Model | | | Longitudinal Model | | |
|---|---|---|---|---|---|---|
| | HR (95% CI) | *p* | $p_{corr}$ | HR (95% CI) | *p* | $p_{corr}$ |
| Global FA | 0.88 (0.32 – 2.41) | .781 | .889 | 0.37 (0.04 – 3.60) | .304 | .359 |
| Anterior Thalamic Radiation | 0.95 (0.36 – 2.48) | .902 | .902 | 0.57 (0.08 – 4.16) | .495 | .511 |
| Cingulum Cingulate Gyrus | 0.80 (0.29 – 2.20) | .636 | .807 | 0.51 (0.14 – 1.83) | .226 | .324 |
| Cingulum Hippocampus | 0.83 (0.26 – 2.71) | .745 | .878 | 0.13 (0.00 – 4.47) | .193 | .317 |
| Corticospinal Tract | 1.06 (0.40 – 2.79) | .900 | .902 | 0.99 (0.31 – 3.20) | .980 | .980 |
| Forceps Major | 1.30 (0.49 – 3.48) | .571 | .797 | 0.28 (0.03 – 2.67) | .201 | .317 |
| Forceps Minor | 0.79 (0.30 – 2.12) | .619 | .807 | 0.47 (0.10 – 2.27) | .265 | .339 |
| Inferior Fronto Occipital Fasciculus | 0.89 (0.33 – 2.44) | .810 | .891 | 0.31 (0.03 – 3.80) | .277 | .339 |
| Inferior Longitudinal Fasciculus | 0.81 (0.28 – 2.29) | .663 | .810 | 0.20 (0.01 – 3.43) | .202 | .317 |
| Superior Longitudinal Fasciculus | 0.76 (0.28 – 2.07) | .561 | .797 | 0.24 (0.03 – 2.05) | .144 | .317 |
| Uncinate Fasciculus | 0.93 (0.35 – 2.48) | .873 | .902 | 0.43 (0.05 – 3.39) | .337 | .370 |

**Table S4**

*Hazard Ratios Consistently <1 Across All White Matter Tracts without Censoring at 3 Years*

| Tract | Baseline Model | | | Longitudinal Model | | |
|---|---|---|---|---|---|---|
| | HR (95% CI) | $p$ | $p_{corr}$ | HR (95% CI) | $p$ | $p_{corr}$ |
| Global FA | 0.58 (0.36 – 0.92) | *.024 | .082 | 0.49 (0.23 – 1.02) | .055 | .116 |
| Anterior Thalamic Radiation | 0.65 (0.41 – 1.01) | .054 | .082 | 0.46 (0.23 – 0.92) | *.033 | .116 |
| Cingulum Cingulate Gyrus | 0.66 (0.42 – 1.06) | .080 | .087 | 0.63 (0.33 – 1.20) | .133 | .146 |
| Cingulum Hippocampus | 0.44 (0.25 – 0.78) | *.007 | .079 | 0.55 (0.23 – 1.33) | .151 | .151 |
| Corticospinal Tract | 0.64 (0.40 – 1.02) | .059 | .082 | 0.57 (0.29 – 1.10) | .081 | .116 |
| Forceps Major | 0.61 (0.39 – 0.96) | *.034 | .082 | 0.50 (0.27 – 0.95) | *.037 | .116 |
| Forceps Minor | 0.70 (0.45 – 1.10) | .116 | .116 | 0.51 (0.28 – 0.94) | *.036 | .116 |
| Inferior Fronto Occipital Fasciculus | 0.63 (0.40 – 0.98) | *.043 | .082 | 0.51 (0.26 – 0.99) | *.048 | .116 |
| Inferior Longitudinal Fasciculus | 0.58 (0.37 – 0.93) | *.026 | .082 | 0.52 (0.24 – 1.13) | .085 | .116 |
| Superior Longitudinal Fasciculus | 0.63 (0.40 – 0.99) | *.046 | .082 | 0.58 (0.29 – 1.16) | .103 | .126 |
| Uncinate Fasciculus | 0.68 (0.43 – 1.05) | .079 | .087 | 0.54 (0.27 – 1.07) | .070 | .116 |

**Figure S1**

*Baseline and Longitudinal Cox Proportional Hazards Models*

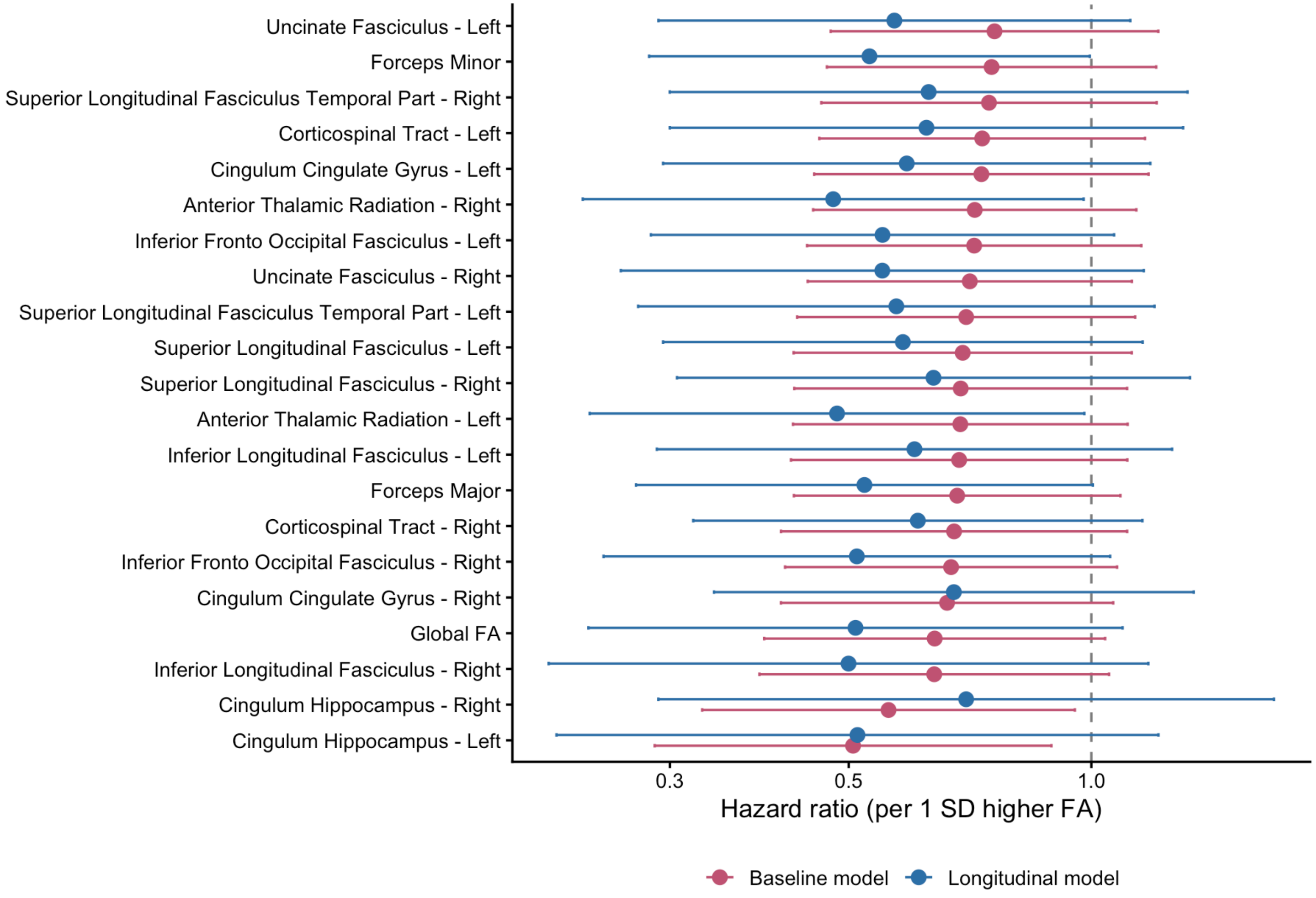
